# A spinal cord injury time and severity consensus transcriptomic reference suite in rat reveals translationally-relevant biomarker genes

**DOI:** 10.1101/2024.01.30.578030

**Authors:** Rubén Grillo-Risco, Marta R. Hidalgo, Beatriz Martínez Rojas, Victoria Moreno-Manzano, Francisco García-García

## Abstract

Spinal cord injury (SCI) is a devastating condition that leads to motor, sensory, and autonomic dysfunction. Current therapeutic options remain limited, emphasizing the need for a comprehensive understanding of the underlying SCI-associated molecular mechanisms. This study characterized distinct SCI phases and severities at the gene and functional levels, focusing on biomarker gene identification. Our approach involved a systematic review, individual transcriptomic analysis, gene meta-analysis, and functional characterization. We compiled a total of fourteen studies with 273 samples, leading to the identification of severity-specific biomarker genes for injury prognosis (e.g., Srpx2, Hoxb8, Acap1, Snai1, and Aadat) and phase-specific genes for the precise classification of the injury profile (e.g., Il6, Fosl1, Cfp, C1qc, Cp). We investigated the potential transferability of severity-associated biomarkers and identified a twelve-gene signature that predicted injury prognosis from human blood samples. We also report the development of MetaSCI-app - an interactive web application designed for researchers - that allows the exploration and visualization of all generated results (https://metasci-cbl.shinyapps.io/metaSCI). Overall, we present a transcriptomic reference and provide a comprehensive framework for assessing SCI considering severity and time perspectives.

**Teaser:** A transcriptomic meta-analysis of spinal cord injury provides a consensus reference and biomarker genes for injury phase/severity.

## Introduction

The manuscript should start with a brief introduction that lays out the problem addressed by the research and describes the paper’s importance. The scientific question being investigated should be described in detail. The introduction should provide sufficient background information to make the article understandable to readers in other disciplines, and provide enough context to ensure that the implications of the experimental findings are clear. USE ACTIVE VOICE: “The team conducted the experiment.” NOT “The experiment was conducted by the team.”

Spinal cord injury (SCI) leads to alterations in the motor, sensory, and autonomic systems immediately below the affected spinal segment, which represent a devastating blow to patients’ quality of life due to the loss of voluntary movement and associated physiological dysfunctions (*1–3*). The World Health Organization (WHO) estimates that 250,000-500,000 people suffer from an SCI each year. Sadly, there is no cure for SCI nowadays, although substantial efforts currently support a wide range of preclinical research aims (*4*, *5*). Recent reports from the Courtine lab have defined a neuronal subpopulation responsible for functional regeneration after epidural electrical stimulation in mice (*6*); nevertheless, we still require concerted efforts in basic research to elucidate the dynamic, complex molecular mechanisms that underlie the distinct stages of SCI. Likewise, an improved understanding of the mechanisms (and their biological relevance) at play at early stages after SCI is needed to control evolution in acute cases and achieve functional recovery in those patients suffering from chronic injuries.

Transcriptomics represents a widely used technology for the study of pathological states such as SCI; this technique provides a comprehensive view of the cellular response to a given condition from a systems biology perspective and allows the identification of differentially expressed genes (DEGs) that may serve as biomarkers for disease diagnosis, prognosis, or provide novel therapeutic targets. Unfortunately, statistically-significant DEGs from studies can display distressingly little overlap when distinct research groups study the same biological system (*7*). This lack of robustness and reproducibility can lead to conflicting or inconclusive results. Although many transcriptomic studies on SCI exist, we still lack a consensus regarding the description of the injury-induced transcriptomic profile and its functional correlation, which hinders the identification of therapeutic targets with sufficient depth over time and based on injury severity due to the dispersion among injuries/models.

To solve such issues, we propose a novel characterization of SCI through a meta-analysis of transcriptomic studies. A meta-analysis represents an integrative approach that allows the measurement of the effect of interest more precisely compared to individual studies, thus increasing statistical power and considering the variability of individual studies to provide more consistent results (*8*). We based our strategy on the systematic review and selection of public transcriptomic studies performed in rats since 2004. We grouped samples based on severity and time after injury, allowing the characterization of SCI in two dimensions. We subsequently processed and analyzed all datasets similarly to avoid introducing biases associated with bioinformatics pipelines (*9*). Our pipeline includes preprocessing, exploratory analysis, and differential gene expression (DGE) analysis. We then performed a gene expression meta-analysis to integrate DGE analysis results, providing a consensus gene expression signature for each group. We identified specific severity- and phase-associated biomarker genes from the generated gene expression patterns. We then functionally characterized these transcriptomic profiles to estimate the dysregulated pathways, infer transcription factor activity, and construct gene co-expression networks. We validated the results bioinformatically with two external datasets and experimentally by quantitative (q-)PCR. We also explored the potential transferability of severity-associated biomarker genes found in rats to predict injury prognosis using human blood samples.

## Results

### Identifying gene consensus signatures based on severity and time

We first performed a systematic review and selection of studies in two public databases for transcriptomics data: GEO (*10*) and ArrayExpress (*11*). A Preferred Reporting Items for Systematic Reviews and Meta-Analyses (PRISMA) flowchart presents the data selection strategy (fig. S1). The search identified 62 unique studies in *Rattus norvegicus*. After applying exclusion criteria, we removed 21 studies since samples did not come from spinal cord tissue or were not analyzed using platforms of interest. Additionally, we also discarded 21 studies as they did not evaluate SCI. After full-text evaluation, we excluded six more studies due to the lack of control samples without treatment or information. Finally, we included 14 studies in our meta-analysis (table S1) (*12–23*).

After discarding samples that received treatment and did not fit the proposed experimental groups, we selected 273 samples from 14 studies for individual analysis. Given the studies diversity, we standardized metadata to create experimental groups with uniform nomenclature. We categorized injury severity into moderate (M) and severe (S) injuries. The severe group included lesions resulting from a spinal cord impact of ≥200 kdyn, a 10 g weight drop from 50 mm, or a complete transection of the spinal cord; meanwhile, we considered injuries not meeting these criteria as moderate (table S2). We established four injury phase groups based on time after injury and sample collection: acute phase (0-3 days, T1), subacute phase (4-14 days, T2), early chronic phase (15-35 days, T3), and late chronic phase (> 35 days, T4) (*3*, *24*, *25*). Controls consisted of samples from sham-operated or naïve rats (non-operated). Figure 1A reports the number of studies per group. Table S3 contains sample identifiers and annotations, while table S4 and fig. S2 provides details on sample distribution into the distinct experimental groups.

**Fig. 1.**
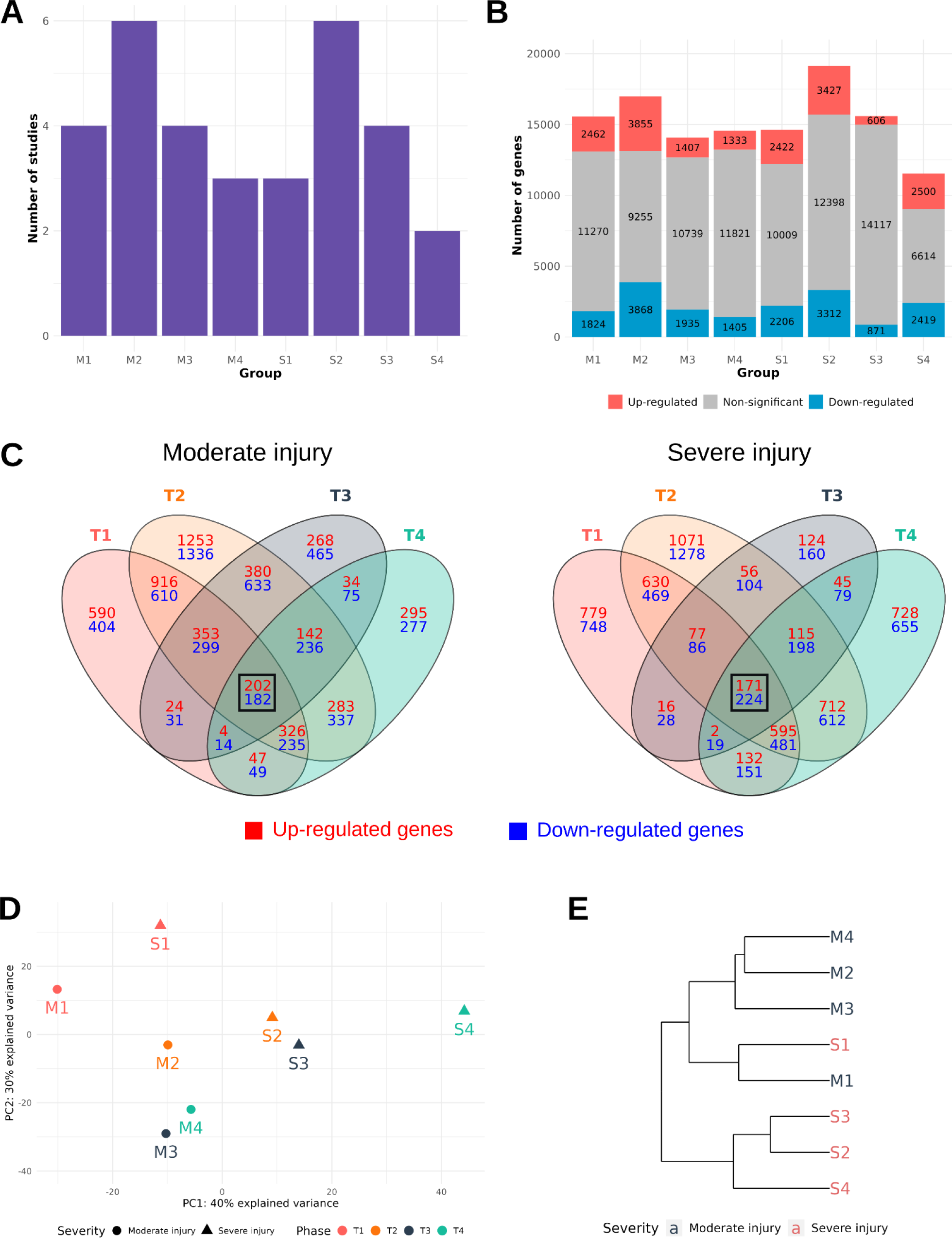
Overview of gene expression meta-analysis results. (**A**) Barplot reporting the number of studies per group. (**B**) Barplot displaying the number of significantly altered genes per group, with significantly up-regulated genes in red, significantly down-regulated genes in blue, and non-significantly altered genes in gray. (**C**) Venn diagrams of significantly altered genes in different injury phases (T1, acute; T2, subacute; T3, early chronic; T4, late chronic) for moderate (left) and severe (right) injury. Intersecting significantly up-regulated genes are noted in red, and significantly dow-nregulated genes are noted in blue. The black square indicates the intersection of all four groups. (**D**) PCA plot and (**E**) hierarchical clustering of logFC gene consensus signatures for all groups.

We followed a common pipeline that included data preprocessing, exploratory analysis, and DGE analysis for the individual analysis of the selected datasets. Since we aimed to characterize SCI in all phases and analyze the influence of injury severity, we compared each injury group against the control group within their study to identify DEGs. The eight comparisons performed and the abbreviations used for simplicity are: M1 (Moderate Acute [T1] vs. Control), M2 (Moderate Subacute [T2] vs. Control), M3 (Moderate Early chronic [T3] vs. Control), M4 (Moderate Late chronic [T4] vs. Control), S1 (Severe Acute [T1] vs. Control), S2 (Severe Subacute [T2] vs. Control), S3 (Severe Early chronic [T3] vs. Control), and S4 (Severe Late chronic [T4] vs. Control). The “Individual analysis” module of the Meta-SCI app contains a detailed list of the individual analysis results. For each dataset and comparison, we indicate the number of DEGs, classified according to their over- or under-expression.

By conducting a meta-analysis, we integrated the DGE analysis results for each comparison to obtain a transcriptional consensus signature for each experimental group. We found an elevated number of significant genes (false discovery rate [FDR] < 0.05) in each comparison in our study (Fig. 1B) and a common core of permanently dysregulated genes at all stages after SCI for both moderate and severe injury (black squares) (Fig. 1C). This result indicates that a considerable proportion of the transcriptional changes occurring after SCI display independence from injury severity and temporal stage. The “Meta-analysis” module of the Meta-SCI app contains a detailed form of the meta-analysis results.

We performed a principal component analysis (PCA) and hierarchical clustering analysis of transcriptional consensus signatures to achieve an overview of the similarities between groups. Together, we observe a separation between severe and moderate injury, with similarities between M1 and S1, suggesting the acute phase as the most distinctive temporal phase irrespective of severity (Fig 1D and E).

### Identification of predictive biomarker genes for severity assessment

To identify severity-specific biomarker genes, we compared transcriptional consensus signatures of the four severe injury groups against the four moderate injury groups (table S5). We identified 282 genes with an FDR < 0.1 (table S6). Interestingly, the hierarchical heatmap revealed a higher gene dysregulation (both up-regulated or down-regulated) in severe compared to moderate injuries (fig. S3), resulting in a clear separation into two distinct heatmap branches based on severity. Fig. 2A and Fig.2B illustrate that the top 10 most significantly altered genes (Srpx2, Hoxb8, Acap1, Snai1, Aadat, Ppic, Lrrc17, Map7, Actg2, and H19) enabled a clear separation between moderate and severe injury, with PC1 explaining 93% of the variance in the PCA. The expression patterns of the top 5 severity-associated biomarker genes demonstrated consistent and more heightened dysregulation over time following severe injury (Fig. 2C), which represents a suitable pattern for injury prognosis.

**Fig. 2.**
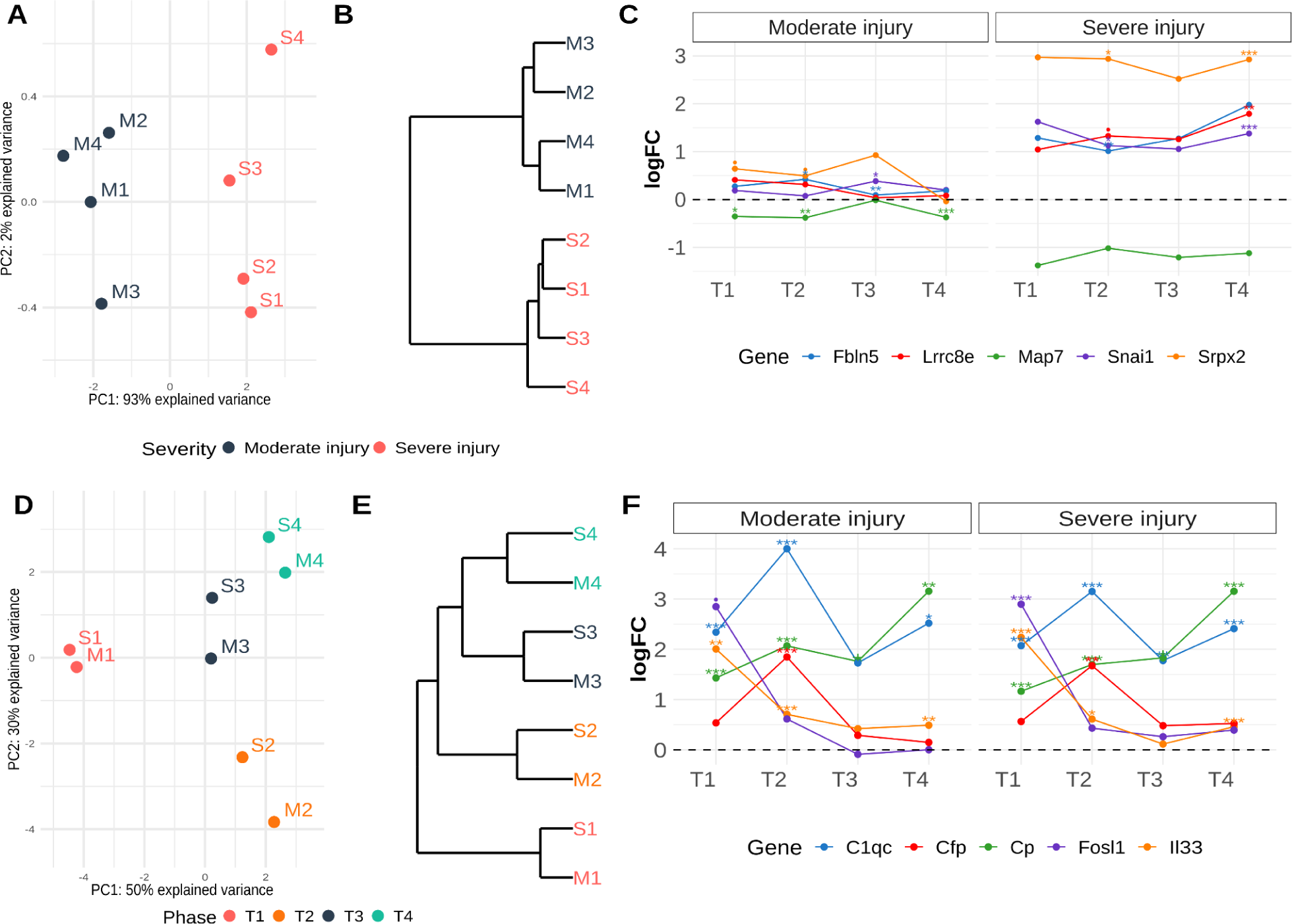
Biomarker genes support effective group classification based on severity or phase after injury. (**A**) PCA reveals a clear separation of groups based on injury severity using the top 10 severity-specific genes, with 93% of the variance explained by PC1. (**B**) The hierarchical clustering of the top 10 genes showcases a clear branching pattern that effectively segregates groups based on injury severity. (**C**) Gene expression patterns of top 5 severity-specific biomarkers. (**D**) PCA plot and (**E**) hierarchical clustering reveal a pairwise temporal classification of groups. (**F**) Gene expression patterns of a selection of different phase-specific genes - Fosl1 and Ill3 as T1-specific; C1qc and Cfp as T2-specific; and Cp as T4-specific.

We divided the list of 282 potential severity-specific genes into up-regulated and down-regulated genes and conducted a protein-protein interaction (PPI) analysis to evaluate functional relationships. The PPI network of down-regulated genes (152) revealed enrichments in functions related to nervous system development and the regulation of trans-synaptic signaling (fig. S4). Meanwhile, the PPI network of up-regulated genes (130) exhibited enrichments in functions related to extracellular matrix organization, animal organ development, and anatomical structure morphogenesis (fig. S5).

### Effective group classification using phase-specific biomarker genes

To systematically identify phase-specific biomarker genes (independent of severity), we compared each phase against the remaining phases. Additionally, we investigated potential phase-specific biomarker genes for each severity (table S5). Table 1 reports the top 10 genes for each comparison. The acute phase displayed the most distinct transcriptional signature, with 770 genes at FDR < 0.1, while other comparisons did not yield significant genes at this threshold. Nevertheless, using the top 10 genes with the lowest p-value in each case allowed clear group separation (fig. S6-9 and table S7-10).

**Table 1.** Top 10 specific biomarker genes for each severity- and time-associated comparison. **Severity**, (S1, S2, S3, S4) vs. (M1, M2, M3, M4); **T1**, (M1, S1) vs. (M2, M3, M4, S2, S3, S4); **T2**, (M2, S2) vs. (M1, M3, M4, S1, S3, S4); **T3**, (M3, S3) vs. (M1, M2, M4, S1, S2, S4); **T4**, (M4, S4) vs. (M1, M2, M3, S1, S3, S4); **M1**, M1 vs. (M2, M3, M4); **M2**, M2 vs. (M1, M3, M4); **M3**, M3 vs. (M1, M2, M4); **M4**, M4 vs. (M1, M2, M3); **S1**, S1 vs. (S2, S3, S4); **S2**, S2 vs. (S1, S3, S4); **S3**, S3 vs. (S1, S2, S4); **S4**, S4 vs. (S1, S2, S3). Genes in bold are upregulated in each comparison, while the others are down-regulated.

| Comparison | Genes |
| --- | --- |
| Severity | **Srpx2**, Hoxb8, **Acap1**, **Snai1**, Aadat, **Ppic**, **Lrrc17**, Map7, **Actg2**, **H19** |
| T1 | **Fosl1**, **Il33**, Vcam1, Mx2, Mdk, **Ldlr**, **Hmgcr**, Alpk3, **Dusp5**, **Serpina3n** |
| T2 | **Cfp**, **Slc46a3**, Scn8a, **Ly86**, **Casr**, **C1qc**, **Slc2a5**, **Fgd2**, **Aif1**, **Slc7a7** |
| T3 | Slc28a2, Iqgap1, Cd14, Nt5dc2, Xdh, Msn, Lyve1, Tmem37, Rdh10, Enpp3 |
| T4 | **Cp**, **Rab27a**, **Rab3b**, **Cybrd1**, **Thrsp**, Plcb1, **Pmp22**, **Acot2**, **Cyp27a1**, **RT1-DI** |
| M1 | **Hmgcs1**, **Fdps**, Cd68, **Gpx2**, Alpk3, **Crym**, **Apln**, Slc15a2, Pltp, **Ldlr** |
| M2 | **Mmp8**, **Cxcl13**, **Cd8a**, **Casr**, **C1qb**, **RT1-CE4**, **Fap**, **Cfp**, **Mmp7**, **Slc7a7** |
| M3 | **Sost**, Rbp1, Cd14, Esm1, Hck, **Plag1**, **Fam102b**, Atf3, **Slc36a3**, **Itgb8** |
| M4 | **Thrsp**, Ccl3, **Tmem204**, **Calml4**, **Cpxm2**, Ckap2, Ttk, Csf3r, **G0s2**, **Mpz** |
| S1 | **Fosl1**, **Il6**, **Kng2**, **Tnfrsf12a**, **Upp1**, **Myc**, **Gprc5a**, C3, **Exo1**, Chodl |
| S2 | **Apoc1**, **Cfp**, **Tnr**, **Slc28a3**, **C1qa**, **Slc46a3**, **Casr**, **Postn**, **Slc40a1**, **Pnlip** |
| S3 | Enpp3, Ccl4, Cd36, Lox, Igsf6, Xdh, Folr2, Msr1, Slc28a2, Rnf4 |
| S4 | **Mmrn1**, **Esm1**, **Anxa8**, **Crabp2**, **Bnc2**, **Igfbp6**, **Slco2a1**, **Ptgis**, **Tspan8**, **Pde5a** |

We selected the top 10 genes from the T1, T2, and T4 groups to create a phase-specific consensus signature, resulting in robust group separation (Fig. 2D and E). In the PCA plot (Fig. 2D), PC1 separated M1/S1 from the remaining groups, while PC2 classified the remaining groups into pairs, with the more similarity between M3/S3 and M4/S4 indicating that the selected biomarkers allow the identification of the injury phase independently of severity. Fig. 2D depicts a similar segregation.

While we identified the top 10 potential biomarker genes for T3 (demonstrating the segregation of M3/S3 from remaining groups), we do not consider them appropriate biomarkers (fig. S8) as their characteristic expression pattern depicts a decrease between T2 and T4. This could be explained by a lack of consensus in results due to study heterogeneity rather than a genuine drop in gene expression during this phase. T3 likely represents a transitional phase between T2 and T4 without any expected expression biological process that peaks in T3, as one might anticipate in T2.

The PPI analysis with upregulated T1-specific genes revealed enrichment in responses to stress, cell cycle, and various metabolic processes (fig. S10). Notably, the up-regulated genes demonstrated an increase in expression after injury that declined over time (as evidenced by gene expression patterns of Fosl1 and Il33 in Fig. 2F, for instance). We also detected genes such as Lcat, Vcam1, or Fxyd1, which become downregulated in T1 but upregulated in the following phases. Downregulated genes displayed enrichments for processes related to nervous system development and lipid metabolism (fig. S11). Overall, these two divergent gene expression profiles represent valuable tools to distinguish T1 from the other stages. By utilizing the top 10 candidate T1-specific genes, PCA1 effectively captures 95% of the variance, resulting in a distinct separation between M1/S1 and the remaining groups (fig. S6). T2-specific genes display gene upregulation peaking in T2 and declining in subsequent phases; the Cfp and C1qc complement genes associated with secondary immune response (23, 24) exemplify this pattern (Fig. 2F). Potential T4-specific biomarker genes displayed an increasing expression pattern from T1 that peaked in T4, as seen for Cp (ceruloplasmin) (Fig. 2F). For T2 and T4, the enrichment of the top 25 genes (arbitrary threshold) again revealed immune response involvement in T2 (fig. S12) and lipid metabolism enrichment in T4 (fig. S13). Overall, our meta-analysis allowed the successful identification of biomarker genes for each injury phase after SCI in independent studies with different rat models.

### Functional characterization of gene consensus signatures

We next performed a functional enrichment analysis for each group via the gene set analysis (GSA) method to characterize the gene consensus signatures obtained in the meta-analysis. Fig. 3A reports the number of significant functions (FDR < 0.05) grouped by database, severity, injury phase, and the direction of enrichment. In all cases, we observed a higher number of up-regulated functions (red) after SCI than down-regulated functions (blue), interpreted as a loss of functionality or down-regulation after SCI. Venn diagrams for moderate and severe injuries indicated a high number of shared functions between the four time phases for up-regulated and dow-nregulated functions (Fig. 3B), suggesting the existence of dysregulated mechanisms induced by the SCI that persist into chronic phases. We used Reactome annotation to classify significantly affected pathways to achieve an overview of the biological categories most affected after SCI. The Reactome database has a hierarchical structure in which biological events at the top include increasingly specific pathways (*26*). This analysis illustrated the frequency distribution of biological events in each comparison for significantly up-regulated (Fig. 3C) and down-regulated (Fig 3D) pathways.

**Fig. 3.**
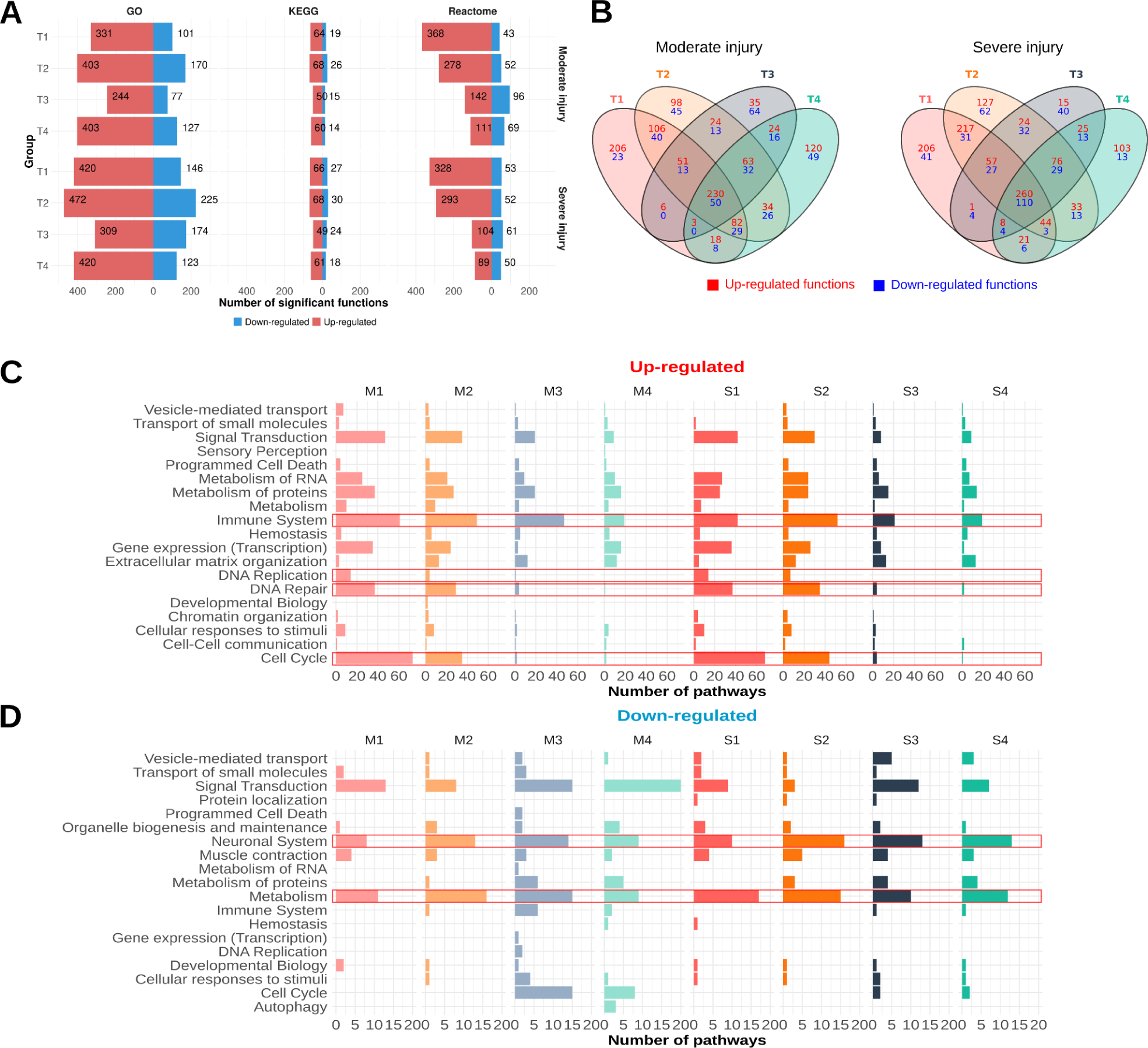
Gene set analysis reveals altered functions/pathways across time and severity after injury. (**A**) Number of significantly altered functions for each database and experimental group. Up-regulated (red) and down-regulated (blue) refer to functions with a logarithm of the odds ratio (LOR) value greater or lower than 0, respectively. (**B**) Venn Diagram of significantly altered functional terms. Relationship between common and unique identified functions in different injury phases (T1, acute; T2, subacute; T3, early chronic; T4, late chronic). Venn diagrams constructed for up-regulated and down-regulated functions in moderate injury (left) and severe injury (right). (**C and D**) Frequency distribution of Reactome categories by phase and injury severity in **C**) up-regulated and **D**) down-regulated pathways. This value represents the sum of all significantly altered pathways within the same biological category in a comparison. Red squares highlight biological categories of interest.

The immune system represents one of the most prevalent categories for significantly up-regulated pathways in all comparisons independently of severity. These pathways form part of the core of permanent dysregulated pathways, which can be explained by the massive infiltration of inflammatory cells during the acute phase, the intrinsic microglial activation (*27–29*) at the injury site, and a failure to efficiently resolve inflammation at the chronic stage (*27*). In contrast, the elevated presence of pathways related to cell cycle, DNA repair, or DNA replication in the acute phase decreases over time, which could be explained by the massive proliferation of spinal cord resident cells such as microglia and macroglia (astrocytes and oligodendrocyte precursor cells) and infiltrated cells such as pericytes, fibroblast or Schwann cells (*23*, *30*). These findings agree with the results described above since T1-specific biomarker genes display enrichments for cell cycle-related functions. The expression pattern of these genes involves a substantial up-regulation in T1 that declines over time, as observed with the number of enriched functions in Fig. 3C (red square).

The permanent loss of neuronal system functions and metabolism characterizes significantly down-regulated pathways. Regarding metabolism-associated functions, the dotplot in fig. S14 reports the down-regulation of functions related to fatty acid synthesis, cholesterol synthesis, and the Krebs cycle in all groups, as previously described (*15*, *31*). Fig. S15 reports in more detail the down-regulation of genes involved in cholesterol synthesis in a hierarchical heatmap, demonstrating the more pronounced level of down-regulation in the chronic compared to the acute phase. Our results agree with the findings of Spann et al. (*15*), who reported the significant down-regulation of the cholesterol biosynthesis pathway and the down-regulated expression of Hmgcs1, Hmgcr, Cyp51, Idi1, and Fdft1, genes involved in the production of cholesterol. We also observed permanent mitochondrial dysfunction, suggested by the down-regulation of functions/pathways related to mitochondria and ATP synthesis (*31*) (fig. S16). The totality of the described events acts in detriment of the correct functioning of the nervous system and any regenerative attempts, as indicated by the large number of down-regulated functions related to synapse regulation, neurotransmitter regulation, or nervous system development in all groups (fig. S17-S19). The “GSA results” module of the Meta-SCI app provides a detailed view of all results; the “Predefined plots” option of the “Dotplots” tool allows an exploration of dotplots of specific related functions; and the “Gene2path” module allows an exploration of the relationship between genes and functions.

### Inference of transcription factors activities dynamics in SCI progression

We next estimated the transcription factor (TF) activity levels based on the expression levels of their target genes, with a methodology that considers whether TF-target interactions activate or repress said target gene (table S11). Thus, the up-regulation (respectively down-regulation) of target genes of an activating (respectively inhibiting) TF becomes interpreted as TF activation following injury [normalized enrichment score (NES) > 0]. Conversely, activating (resp. inhibiting) TFs with down-regulated (resp. up-regulated) target genes, are interpreted as lost TF function or deactivation following injury (NES < 0).

The assessment of TF activity across the different experimental groups revealed a generally higher prevalence of activated vs. deactivated TFs in all scenarios (Fig. 4A). These findings underscore the more substantial impact on TF activation during the early stages after SCI, which gradually decreases in the chronic phases. The Venn diagrams in Fig. 4B reveal 19 and 40 TFs with common differential activation patterns across all phases in moderate and severe injuries, respectively.

**Fig. 4.**
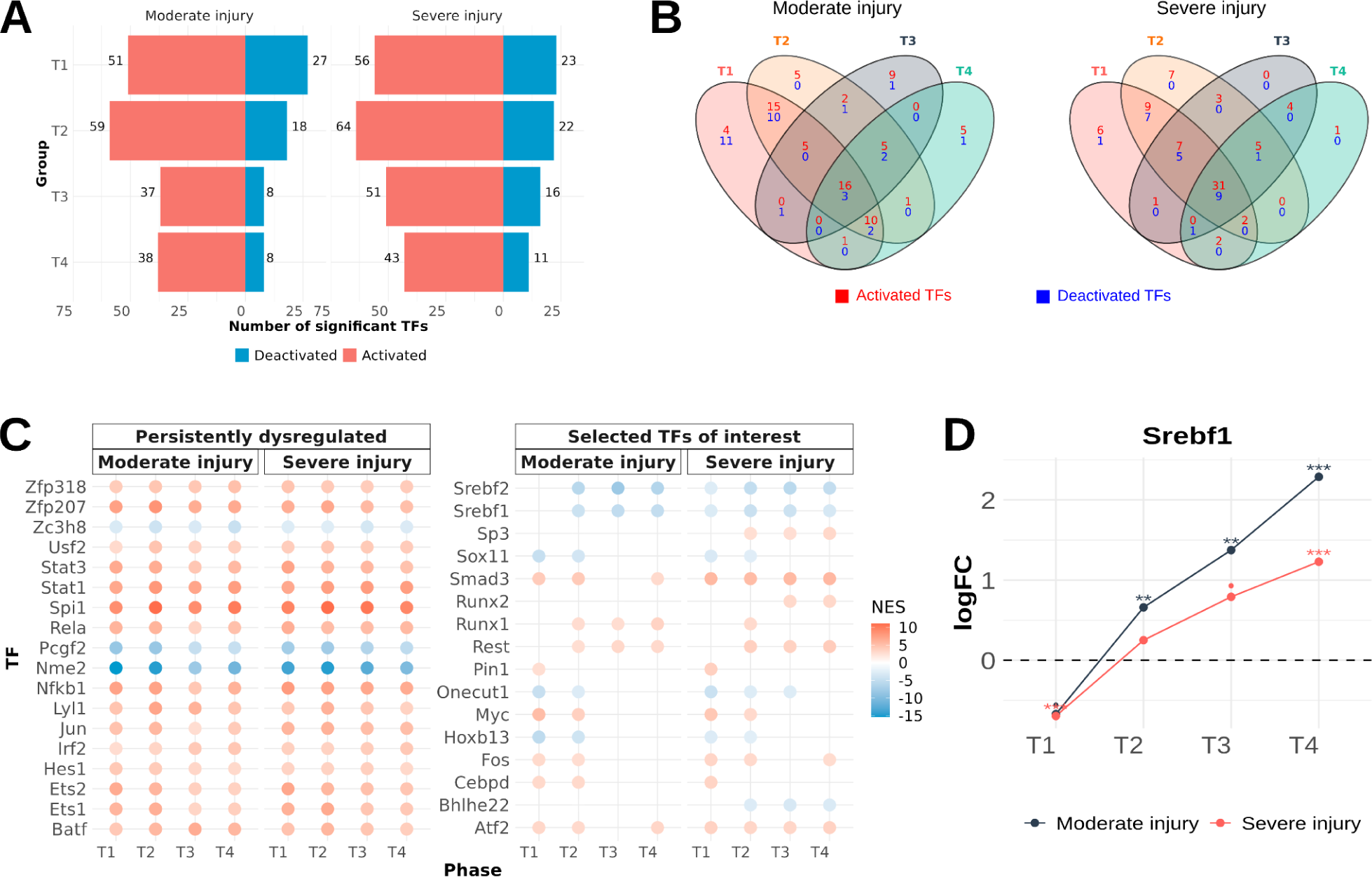
Summary of transcription factor activity inference analysis. (**A**) Number of significantly altered TFs for each experimental group. Activated (red) and deactivated (blue) refer to TFs with a NES value greater or lower than 0, respectively. (**B**) Venn Diagram of significantly altered TFs. Relationship between the common and unique identified functions in different injury phases (T1, acute; T2, subacute, T3., early chronic, T4., late chronic). Venn diagrams constructed for up-regulated and down-regulated functions for moderate (left) and severe injury (right). (**C**) Dotplot of significant dysregulated TFs in all groups (left) and selected TFs of interest with significantly different activities (right) (Red – activation; blue – deactivation).

Fig. 4C (left panel) depicts consistently dysregulated TFs shared between moderate and severe injury. TFs such as Nfkb1, Stat1, and Stat3 become activated during injury progression, contributing to processes such as inflammatory response modulation (*32–34*). We also identified previously undescribed differentially-activated TFs associated with SCI (e.g., Zfp207, Zfp318, and Zc3h8) that open new avenues for the design of therapeutic strategies.

The right panel of Fig. 4C illustrates the activity inference for selected severity- or phase-associated TFs. Sp3, Runx2, and Bhlhe22 exhibited distinct activity profiles in severe compared to moderate injury; meanwhile, TFs such as Pin1, Myc, Cebpd (differentially-activated), and Hoxb13 and Sox11 (differentially-deactivated) undergo deregulation during acute phases but revert to normal levels following development into the chronic phase. Interestingly, Srebf1 exhibits an increasing up-regulated gene expression pattern (Fig. 4D); however, this TF becomes differentially-deactivated in all comparisons except M1. Both Srebf1 and Srebf2 (also deactivated) represent activators of lipid metabolism genes (*35*), which could explain the previously described down-regulation of fatty acid and cholesterol synthesis.

### Functional classification of gene co-expression networks

We use the gene consensus signatures matrix (table S16) from moderate and severe injuries separately as input to find genes with similar expression patterns. We identified 19 and 18 clusters of co-expressed genes in moderate and severe injuries, respectively, and constructed a PPI network for each cluster. The extensive constructed networks displayed high levels of interconnection, indicating the functional association of genes with similar expression patterns. To gain deeper insights, we generated 261 subnetworks, conducted a functional enrichment analysis for each subnetwork, and then classified them into manually predefined biological categories of interest. Among the 261 generated subnetworks, we successfully classified 153 (Fig. 5A). The remaining 108 subnetworks comprised only a few proteins without enriched functions. The “PPI networks” module of the Meta-SCI app enables in-depth exploration of all generated networks.

**Fig. 5.**
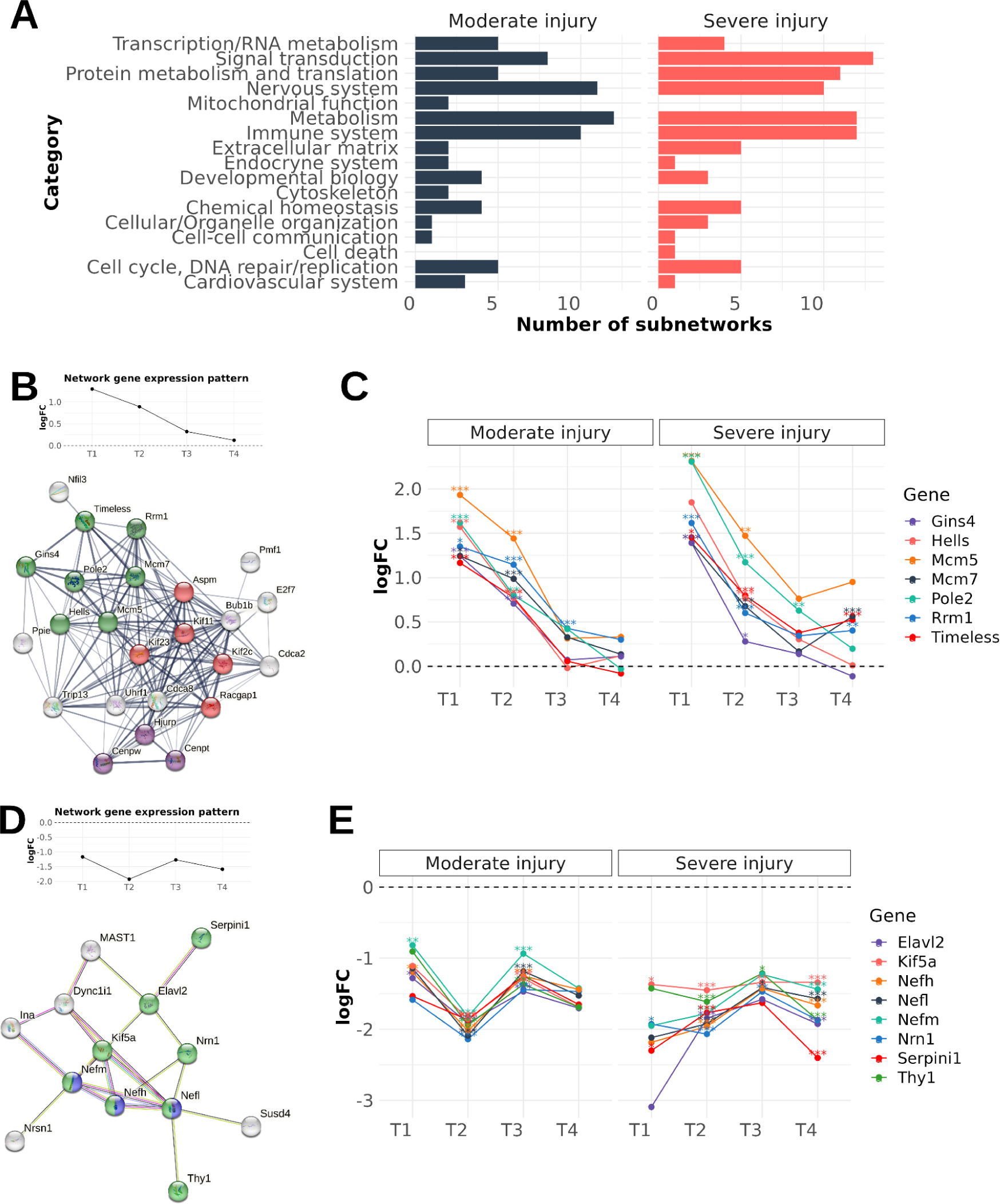
Summary of co-expression network results. (**A**) Frequency distribution of functional categories among networks. (**B**) Moderate injury - cluster 11, subnetwork 1 - functionally relates to the cell cycle. All subnetwork proteins display a functional relationship and exhibit the same expression pattern. Moreover, red, green, and purple proteins form physical complexes among themselves. (**C**) Gene expression patterns of green proteins in moderate injury cluster 11, subnetwork 1. (**D**) Moderate injury - cluster 6, subnetwork 1 - functionally relates to the nervous system. All subnetwork proteins display a functional relationship and exhibit the same expression pattern. Proteins annotated to “generation of neurons” (GO:0048699) are shown in green, and “spinal cord development” (GO:0021510) is shown in blue. (**E**) Gene expression patterns of green proteins in moderate injury - cluster 6 subnetwork 1.

As observed in the GSA results, we found the prominent representation of categories such as nervous system, immune system, and metabolism; moreover, we observed notable functional similarity across severity levels. Fig. 5B depicts cluster 11, subnetwork 1 for moderate injury, comprising genes related to the cell cycle (proteins in green, red, and purple also form a physical part of the same complexes). These genes possess an expression pattern characterized by an initial upregulation that decreases over time (Fig. 5C); as discussed previously, this explains the presence of enriched functions related to the cell cycle in T1 that decrease in subsequent phases. Fig. 5D displays cluster 6, subnetwork 1 for moderate injury, comprising genes involved in the nervous system. Genes associated with the “generation of neurons” (GO:0048699) are shown in green, while those involved in “spinal cord development” (GO:0021510) are shown in blue; in this case, a consistent downregulation of genes characterize this expression pattern (Fig. 5E).

### Consistency in temporal gene expression patterns between the mouse and rat

To validate and compare gene expression patterns obtained in our meta-analysis, we selected the mouse RNA-seq dataset from Li et al., 2022 (*36*), which possessed an experimental design similar to the studies included in our work. The authors used a mouse model to study gene expression changes following SCI. They induced injuries at the T9 level and collected samples from a spinal cord segment at different days post-injury (dpi) that covered the four times phases established in our work. We then followed the same pipeline applied to individual studies to generate gene expression patterns, ensuring their comparability with those generated in our meta-analysis. Table S12-S16 reports the DGE analysis results of each comparison and the gene expression patterns matrix.

We then conducted a correlation analysis to compare gene expression patterns between the mouse and rat models (rat moderate injury vs. mouse; rat severe injury vs. mouse; rat moderate injury vs. rat severe injury), which demonstrated a generally positive relationship between them. The density plot of correlation values indicates a median close to 0.5 in three comparisons (Fig. 6A and table S17). A total of 892 genes displayed a strong positive relationship (r > 0.9) between mouse and moderate rat injury, another 892 genes demonstrated a positive relationship between mouse and severe rat injury, and 922 genes presented a positive relationship between moderate rat injury and severe rat injury. Among these, 198 genes exhibited a positive correlation across all three comparisons (Fig. 6A). We next performed a clustering analysis using a curated set of potential phase-specific genes (Fosl1, Il6, Cfp, Sipa1, Cp, and Dbp) that display similar gene expression patterns across the three comparisons (Fig. 6B). Similar to Fig. 2C and Fig. 2D, we observed correct stratification into temporal phases. Hierarchical clustering revealed a clear separation of groups based on time post-injury, where T1 lies isolated in an independent branch, followed by the segregation of T2 from the chronic phases (T3 and T4). In this case, the mouse T3 and T4 signatures become grouped within the same branch as M3 and S3 of the rat, while M4 and S4 remain independent (Fig 6C). These findings could be explained due to slight differences in timings - mouse T3 is defined as 28 dpi, and T4 as 42 dpi in the mouse dataset, which lie nearer in time when compared to T3 and T4 from individual studies of rat meta-analysis. Rat T3 covers 15 to 35 dpi, and T4 extends from 56 to 168 dpi. Longer intervals between phases could lead to more distinct differences in expression profiles, as the injury has additional time to stabilize.

**Fig. 6.**
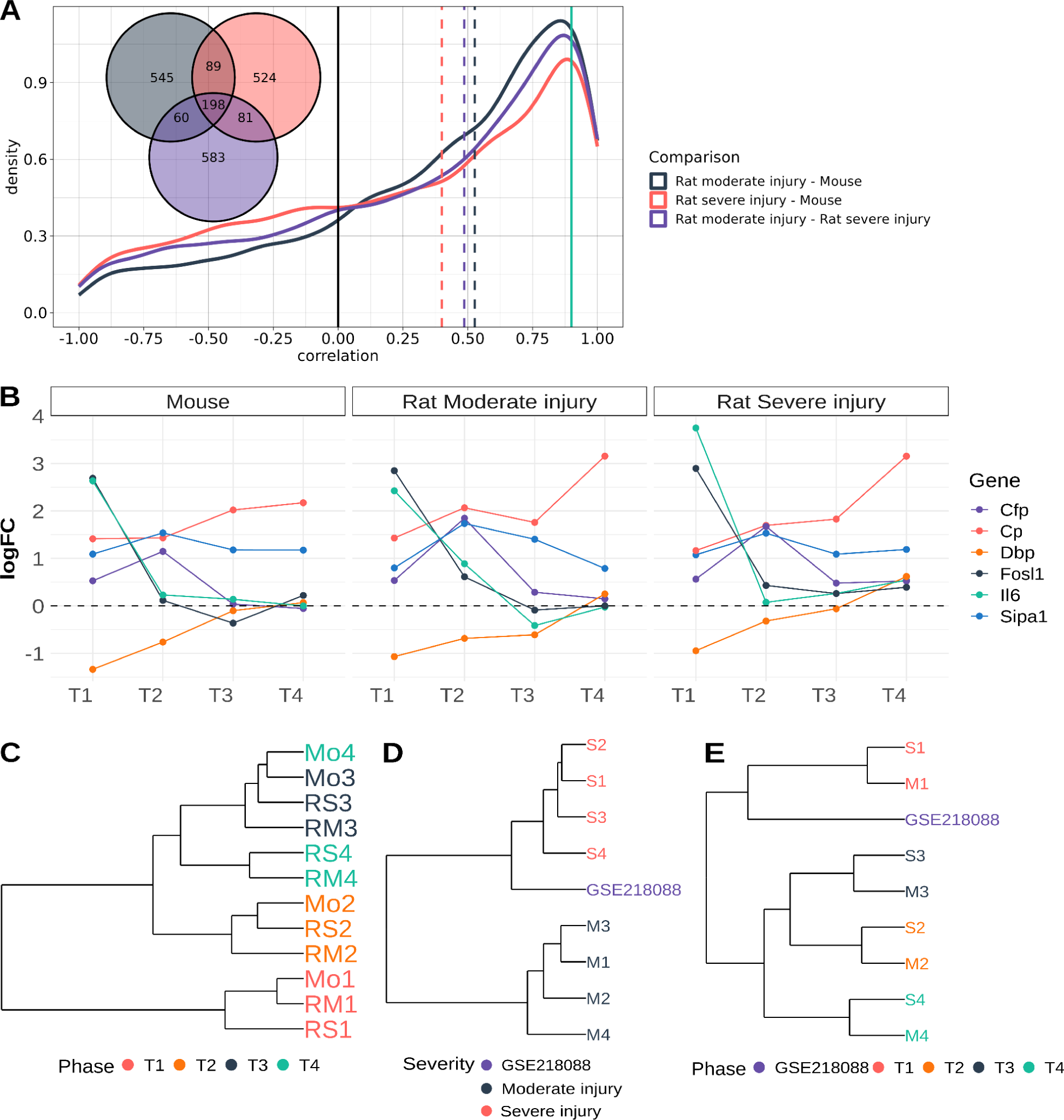
Gene expression patterns comparison between rat meta-analysis and mouse dataset, and clustering analysis using biomarker genes in mouse and GSE218088 datasets. (**A**) Density plot of correlation values between gene expression patterns. Black denotes rat moderate injury vs. mouse; red denotes rat severe injury vs. mouse; and purple denotes rat moderate injury vs. rat severe injury. The green line represents a correlation coefficient of 0.9. Dashed lines represent the median correlation values for each comparison. The Venn diagram depicts the intersection of genes with a correlation coefficient greater than 0.9 in each comparison. (**B**) Comparison of expression patterns of phase-specific genes in mice and moderate and severe injury in rats. (**C**) Hierarchical clustering using phase-specific genes (Fosl1, Il6, Cfp, Sipa1, Cp, and Dbp) with meta-analysis signatures and mouse dataset. (**D**) Hierarchical clustering using severity-specific genes with meta-analysis signatures and GSE218088 transcriptional signature. (**E**) Hierarchical using phase-specific genes with meta-analysis signatures and GSE218088 transcriptional signatures.

Our comparative analysis with an SCI mouse model validated the gene expression patterns established in our meta-analysis and demonstrated a positive correlation between mouse and rat injury responses, supporting the reliability of the identified biomarker genes.

### Our proposed severity- and phase-specific biomarker genes effectively classify novel rat transcriptomic profiles

We used the GSE218088 dataset (*37*) published after the study search and selection period as a validation set, given that it meets the inclusion criteria established in the meta-analysis. We aimed to evaluate whether the selected phase- and severity-specific biomarker genes could classify a new transcriptomic profile correctly. In this new study, the authors indicated that the free fall of a 10 g hammer with a diameter of 2.5 mm produced the injury, which is considered severe (*38*, *39*). The authors indicated that they extracted spinal cord tissue three days after SCI, corresponding to the acute phase (T1) defined in our study.

Using the top 10 severity-specific genes (Table 1), we observed the clustering of the transcriptional signature of the GSE218088 dataset with the severe injury groups in our analysis (Fig. 6D). Additionally, using phase-specific biomarkers for T1, T2, and T4 (Table 1), we observed the clustering of the transcriptional profile of GSE218088 with the T1 signatures from our meta-analysis (Fig 6E). Therefore, this comparison demonstrates the robustness of the results obtained and the capacity of the biomarker genes to classify a new transcriptional profile.

### Concordance between meta-analysis gene expression patterns and qPCR analysis

We aimed to confirm through experimental validation the expression patterns identified in our meta-analysis. Following severe injury induction, we extracted spinal cord RNA from injured rats at T1, T2, T3, and T4 post-injury. We selected eight genes to validate their expression profiles by qPCR. We chose four TFs (Cebpd, Hes1, Jun and Onecut1) due to their functional relevance, whereas the other four genes (Il6, Vcam1, Vegfa, and Vim) represent targets of these TFs and potential phase-specific biomarker genes. While reproducing the exact expression pattern remained challenging, this validation confirmed general dysregulation patterns or at least the direction of dysregulation at specific phases after injury (Fig. 7).

**Fig. 7.**
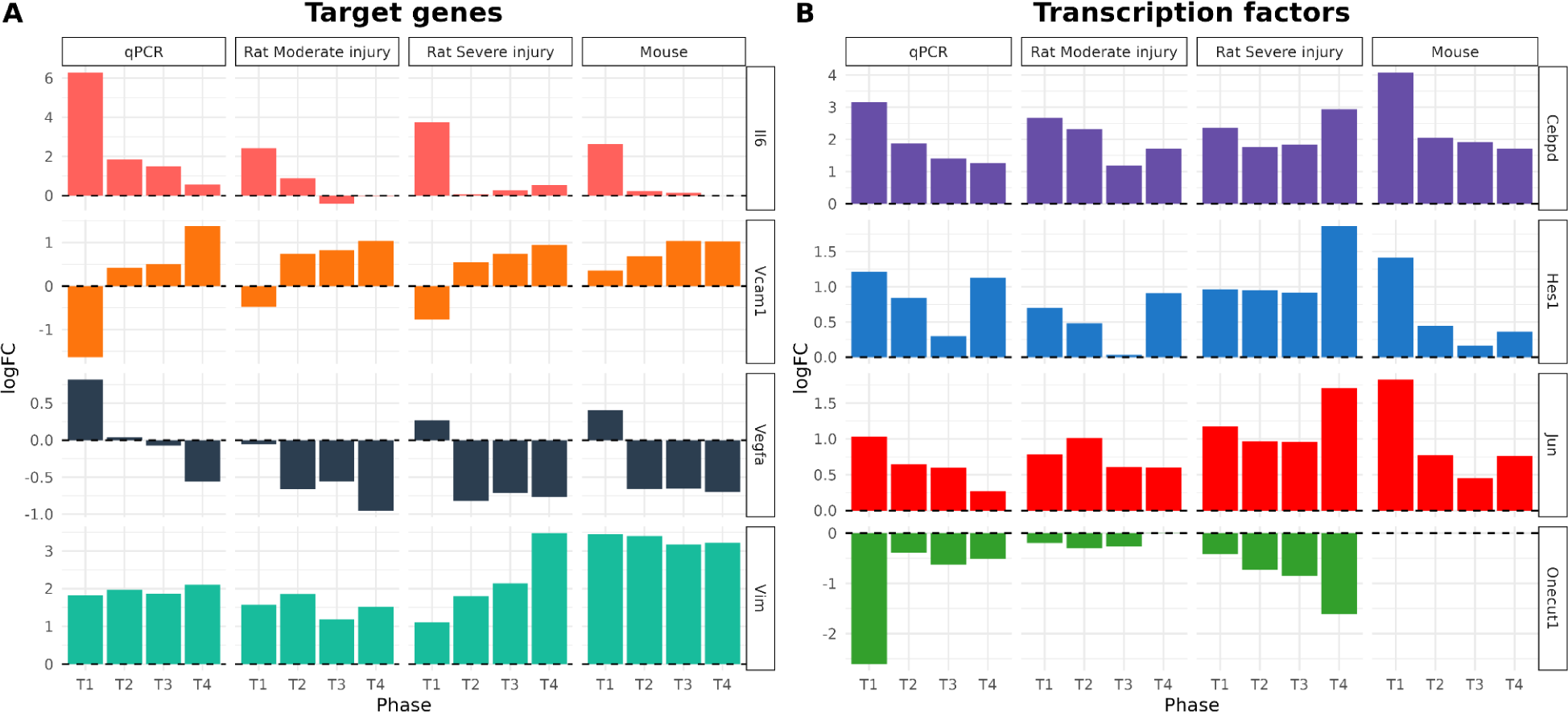
Gene expression patterns comparison of selected TFs and targets for experimental validation. qPCR values were obtained by calculating the logFC of the expression at each time vs. the expression of the uninjured samples in qPCR. Rat Moderate injury indicates the expression pattern obtained in the meta-analysis for moderate injuries. Rat Severe injury indicates the expression pattern obtained in the meta-analysis for severe injuries. Mouse indicates the expression pattern obtained in the mice dataset.

In particular, Il6 exhibited general overexpression with notably higher expression in T1 than in other phases, in which it remained similar to pre-injury levels. Vcam1 presented down-regulation in T1 followed by a gradual up-regulation expression increase over time, except in the mouse dataset, where we only appreciate the gradual expression increase over time. Oppositely, Vegfa showed up-regulation in T1 and down-regulation in the other phases. Vim confirmed the constant up-regulation, although qPCR did not validate its potential role as S4-specific biomarker.

With respect to TFs, Cebpd, Hes1 and Jun presented similar up-regulated patterns in qPCR, meta-analysis and mouse dataset. Although there is lower consistency in the expression patterns across all four systems, we observe a general agreement. Contrarily, Onecut1 presented different patterns of change but confirmed its constant down-regulation over time.

### Severity-specific rat biomarker genes predict injury prognosis from blood samples from human SCI patients

To investigate the potential clinical translation of severity-specific biomarker genes identified in rats to the prediction of injury prognosis from blood samples of human SCI patients, we analyzed the GSE151371 dataset (*40*), which contains gene expression profiles of human blood samples collected in the first 10 days after SCI. Human SCIs are categorized based on severity and prognosis using the American Spinal Cord Injury Association (ASIA) with a 5-grade scale- the American Impairment Scale (AIS) - ranging from A-E, moving from more to less severe. Grade A denotes complete sensory and motor loss; Grade B signifies complete motor loss but preserved sensation; Grade C and D represent various degrees of motor function preservation; while Grade E indicates normal sensory and motor function (*41*).

To achieve a similar scenario to our rat-based meta-analysis, we selected samples from AIS A (complete sensory and motor loss) and AIS D (a certain degree of functional preservation) (*41*) to ensure similarity in severity between the two biological systems. After DGE analysis between groups, we identified 931 genes (FDR < 0.1; table S18); we compared this list with the 282 severity-specific genes previously identified in rat models (Fig. 8A). We found 12 intersecting genes (EXT1, FBN1, FOSB, GNAO1, GRB10, MMP9, NFE2, PRF1, SLC25A23, SNAI1, ST6GALNAC3 and STX11), which expression pattern in the GSE151371 dataset is characterized by an increasing dysregulation as SCI severity increases (fig. S20).

**Fig. 8.**
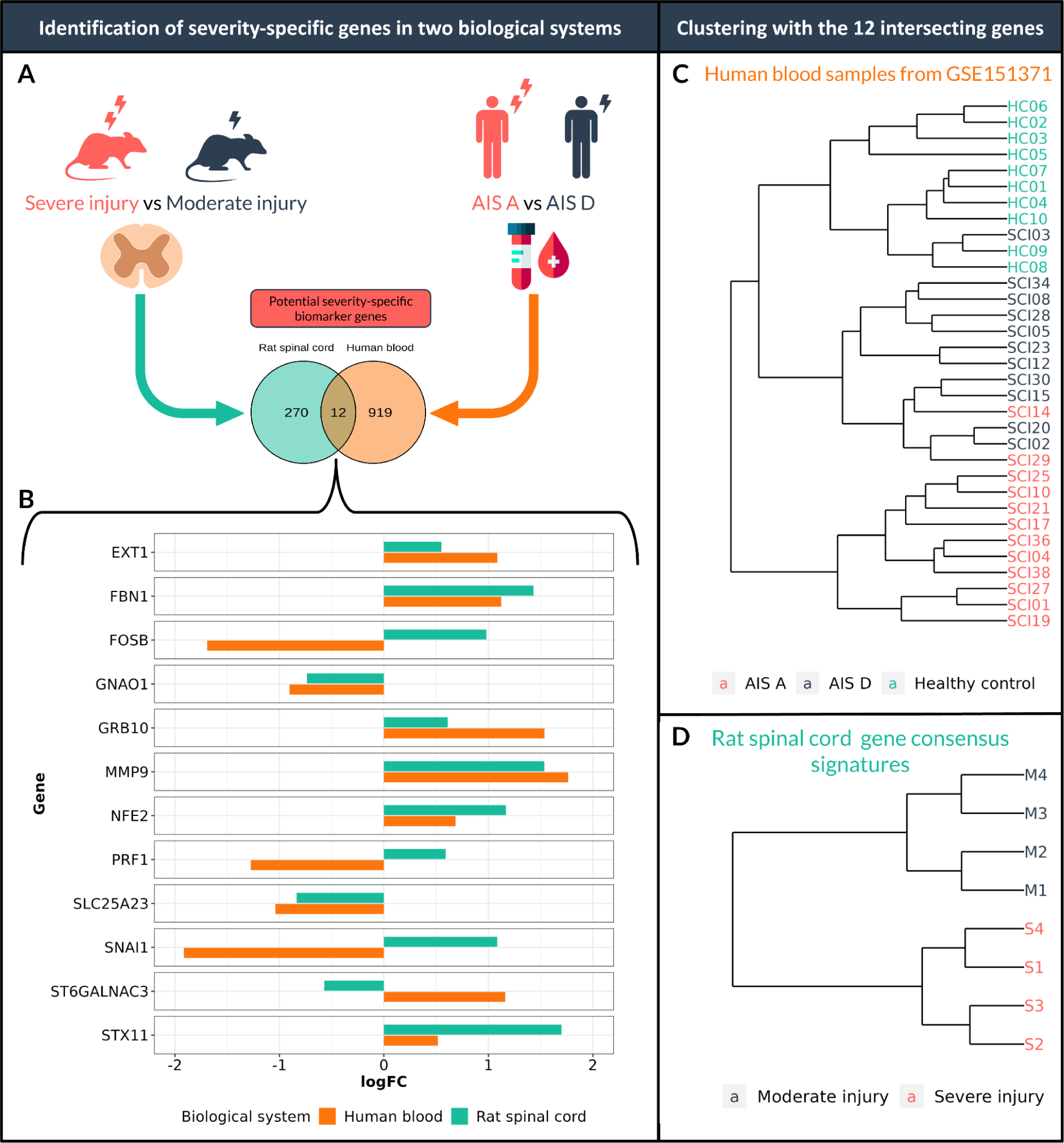
Severity-specific biomarker genes stratify samples in human and rat SCI models. (**A**) Schematic illustration of the approach to identify severity-specific biomarker genes common to the two biological systems. **(B)** Barplot comparing the logFC of the 12 intersecting genes. Orange bars indicate the logFC values after comparing AIS A vs. AIS D in human samples. Green bars indicate the logFC after comparing the consensus logFC in severe injuries and consensus logFC in moderate injuries obtained in rat meta-analysis. **(C)** Hierarchical clustering uses the 12-gene signature in human blood samples from the GSE151371 dataset. **(D)** Hierarchical clustering using the 12-gene signature in rat spinal cord gene consensus signatures obtained in the meta-analysis.

We then used this 12-gene signature to analyze its ability to stratify groups according to severity by clustering analysis in rats and humans. Human blood sample analysis revealed the presence of three clusters: severe injury, moderate injury, and healthy controls (Fig. 8C), while the gene consensus signatures demonstrated stratification based on the severity level in rats (Fig. 8D). These findings indicate that expression levels of the 12 identified biomarker genes can delineate severity-specific subgroups in human blood and rat spinal cord samples.

### Meta-SCI app - an interactive, user-friendly platform to explore results

We have developed the Meta-SCI app (https://metasci-cbl.shinyapps.io/metaSCI/), an interactive web application that provides a user-friendly interface to consult and visualize the results obtained in this work, allowing users to gain a deeper understanding of the underlying mechanisms of SCI.

This application consists of eight main modules (Fig. 9). The “Meta-analysis” module enables the generation of forest and funnel plots for genes of interest and provides access to all meta-analysis statistics. The “Clustering” tool facilitates PCA and clustering analysis of consensus profiles using user-selected genes or predefined sets, aiming to identify group classification related to gene expression patterns over time and severity. Clustering analysis can be performed on all eight consensus profiles or just one severity model; additionally, functional clustering can be conducted based on the GSA results. The “Gene patterns” section compares the time and severity of transcriptional changes in genes of interest and also helps identify highly correlated gene patterns between moderate and severe injuries. The “TF activities” section offers hierarchical heatmap visualization based on the expression of TF target genes and also provides comprehensive statistics for TF activity inference and individual gene targets. Within the “PPI networks” module, users can explore networks constructed by co-expressed genes and their functional roles. All the available STRING web analyses can be used directly in this module. Users can search for networks containing a gene of interest or networks related to a specific functional category or functional term. The “GSA” module comprises 3 different tools. With “GSA results,” users can access all GSA statistics in tabular format. Using the “Dotplot” tool, users can generate dot plots for functions/pathways of interest, search them by keywords, or visualize predefined graphs of relevant biological categories. All plots are dynamically reactive to the applied filters. The “gene2path” section is similar to the TF activities tool, but the hierarchical heatmaps display the expression level of genes belonging to a functional term, and tables show the GSA stats. The “Validation set” module integrates the previously mentioned tools, Gene patterns, and Clustering. In this instance, the module facilitates the comparison of expression patterns with an external mouse dataset and performs clustering analysis of mouse and rat transcriptomic profiles, enabling the identification of common biomarkers in both species. Finally, the “Classify your data” tool enables users to analyze their expression profiles and evaluate their similarity to the meta-analysis consensus profiles based on user-selected or predefined biomarker genes.

**Fig. 9.**
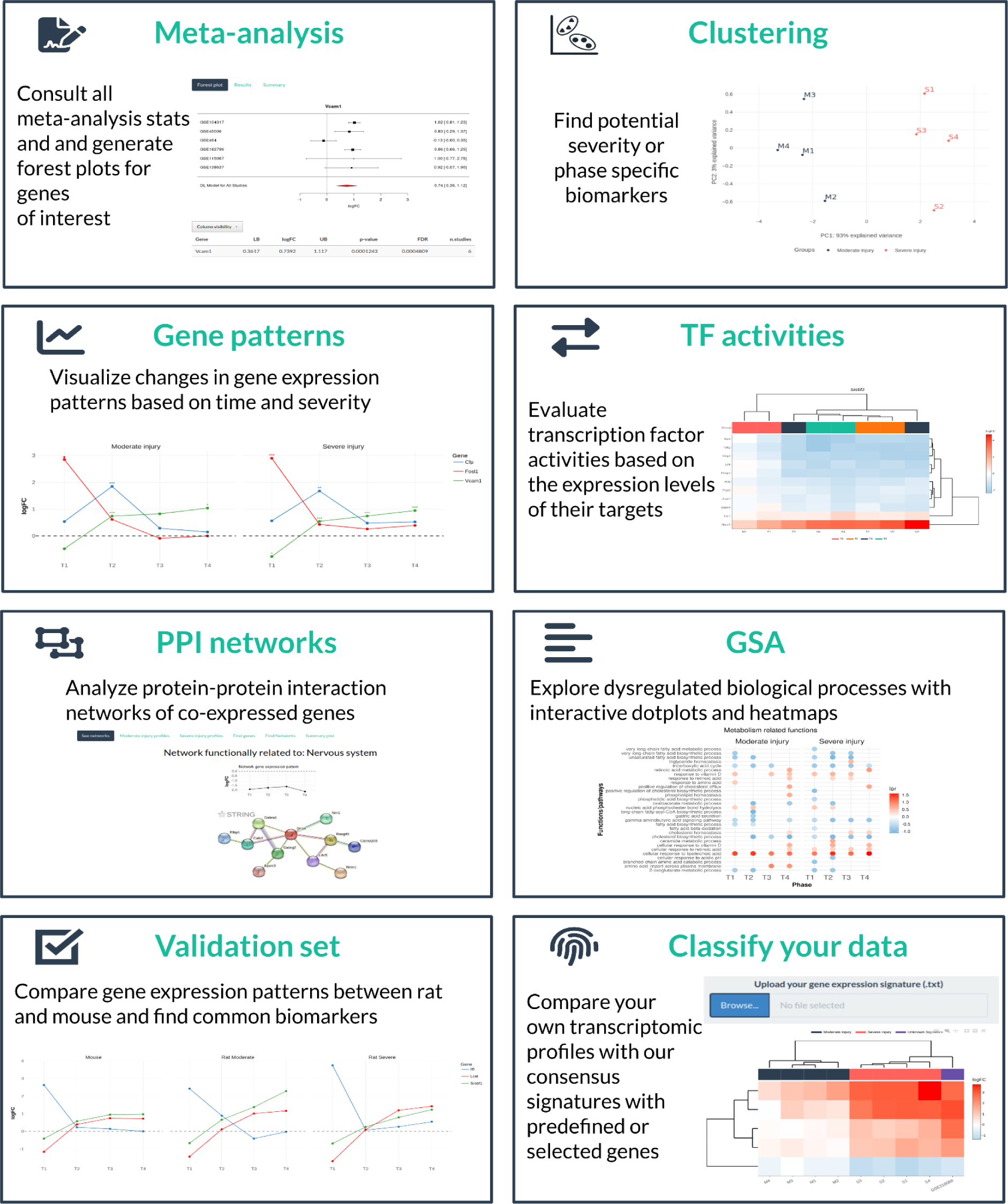
Overview of the main modules of the Meta-SCI app. Meta-SCI app has eight main modules: 1) Meta-analysis; 2) Clustering; 3) Gene patterns; 4) TF (transcription factors) activities; 5) PPI (protein-protein interaction) networks; 6) GSA (gene set analysis); 7) Validation set; 8) classify your data. Meta-SCI app is available at https://metasci-cbl.shinyapps.io/metaSCI/.

The Meta-SCI app is an innovative tool for researchers interested in SCI. All plots and data presented in the app are dynamic, allowing users to interact with them and explore the results in more detail. Additionally, users can download the data and plots, enabling further analysis and research. For additional information on the modules of the Meta-SCI app, consult the “Help” section.

## Discussion

The inherent complexity of SCI has prompted the ongoing and urgent development of novel approaches to enhance the understanding of underlying disease-associated mechanisms. Here, we present a transcriptomic meta-analysis consolidating the most significant number of studies/samples regarding rat SCI datasets to our knowledge. Furthermore, we integrated diverse experimental models based on severity and phase post-injury. As a pioneering effort, we also established a freely accessible transcriptomic reference for SCI research.

The strength of our work emerges from the generation of transcriptomic consensus signatures within established groups. The application of uniform processing and integration through meta-analysis enhances the statistical robustness in determining the magnitude of gene alteration, mitigating inherent dataset heterogeneity and biases stemming from disparate pipelines – a significant feat, given that studies with identical experimental designs can yield contradictory expression changes for the same gene (*42*). Experimental validation represents another strength of our study, which we demonstrated via i) qPCR of eight gene signatures, confirming in all cases the direction of dysregulation; ii) bioinformatic analysis, finding a positive correlation in temporal transcriptional changes between rat and mouse; and iii) the correct classification of an external transcriptional signature using the proposed severity and phase biomarker genes. Our work facilitates the comparison of gene expression levels, pathways and TF activity across time/injury severities. Furthermore, direct identification of gene expression levels associated with specific pathways or TF targets is possible. All results are dynamically and interactively accessible through the Meta-SCI app. While one of our study’s strengths lies in assembling 14 studies and 273 samples, unequal representation across experimental groups represents the main limitation, highlighting an emphasis on acute and a lack of samples in chronic phases. For example, the M2 and S2 groups include samples from six studies, while the S4 group contains samples from only two. As far as our knowledge extends, Squier et al. previously integrated the larger number of rat SCI transcriptomic datasets. Nevertheless, our research represents progress as we incorporated a more comprehensive array of diverse datasets (14 vs. 3) and employed more sophisticated integration methods using meta-analysis instead of a basic intersection of results. This underscores the necessity to publish data following FAIR principles (findability, accessibility, interoperability, and reusability) (*43*), as certain studies do not provide data or adequate sample descriptions. Another potential limitation arises from including datasets derived from diverse technologies and platforms (10 microarrays and 4 RNA-seq experiments). However, the standardized reanalysis and subsequent integration after DGE analysis effectively captured changes in transcriptional regulation (*9*).

This study generated eight consensus transcriptional profiles that comprehensively characterized each predefined group. These transcriptional profiles serve a dual purpose: first, they provide insight into phase- and severity-specific expression patterns of genes, and second, they were used for the identification of potential biomarkers and functional characterization. The robustness of these expression profiles is reinforced by different sources. Previously, Tica et al. (*44*) combined four transcriptomic and proteomic datasets to identify genes/proteins persistently dysregulated at 7 days and 8 weeks post injury. They found 40 up-regulated and 48 down-regulated genes/proteins, which consistently matched our meta-analysis results. Moreover, these genes exhibited persistent dysregulation in moderate and severe injuries. Notably, our research identified one of these genes - Vim - as consistently up-regulated across all time points after SCI, corroborating results from the meta-analysis, the mouse dataset, and qPCR validation. We also validated the expression profiles of the meta-analysis by comparing them with the expression profiles of a mouse dataset (*36*), which shared a very similar experimental design with the studies included in our work. We observed an overall positive correlation between mouse and rat gene expression patterns for moderate and severe injury. When comparing these three systems, 2080 unique genes exhibited correlation coefficients greater than 0.9 in at least one of the three comparisons, with 198 genes intersecting across all biological systems, which reinforces the concept of model similarity and the potential translatability of discoveries. While the exact replication of expression patterns posed challenges, qPCR-based validation reaffirmed the direction of dysregulation (up-regulation or down-regulation) at specific time points. The qPCR validation underscored the robustness of our meta-analysis-derived expression patterns. Overall, these transcriptomic consensus signatures provide insight into how gene expression patterns evolve, offering valuable information regarding the development of potential therapeutic strategies suitable for each SCI phase.

We successfully identified biomarker genes that clearly distinguish moderate and severe injury, displaying significant and persistent dysregulation in severe injuries. We examined whether the 282 potential severity-specific genes displayed functional interconnection through PPI analysis. The down-regulated genes formed a highly connected network, playing roles in the functions and development of the nervous system and synapses. These findings mirror the increased loss of function associated with severe injury, given the greater extent of damaged tissue and complete axonal rupture. The up-regulated genes created a tightly interconnected network associated with tissue remodeling and extracellular matrix organization. Notably, 45 of these genes are translated into proteins that will be located in the extracellular region. In severe injuries affecting more prominent areas and suffering more damage, these genes might become activated to regulate mechanisms involved in wound healing and fibrosis (*45*). By employing the top 10 severity-specific genes, we effectively classified an external dataset involving a severe rat injury, demonstrating their validity. The expression profiles of the identified severity-specific genes make them suitable as potential prognostic biomarkers. Since no significant expression changes occurred in moderate damage, but in severe injuries there is considerable and persistent dysregulation starting from T1, these genes can be employed at the early stages of injury to predict its progression. Among potential severity biomarkers, we identified matrix metalloproteinases (MMPs), Mmp9 and Mmp23, which exhibited significantly higher up-regulation in severe compared to moderate injury. Previous studies suggested MMPs as potential predictive biomarkers for worse neurological outcomes (*46–50*). Kwon et al. (*48*) described a significant correlation between elevated MMP9 levels and worsened neurological recovery in human patients. Interestingly, studies conducted in canine models observed higher MMP9 levels and a poorer prognostic profile in animals with more severe SCI (*50*). These findings suggest that the same prognostic biomarkers for injury outcomes could be identified in animal models and patients, reinforcing our results’ validity and potential translation to clinical research. In addition, Microtubule-Associated Protein 2 (Map2) has previously been proposed as a severity biomarker (*47*, *51*). Our results indicate a greater down-regulation of the expression of Map2 (and Map7) in severe injury, which could represent a new candidate biomarker gene for future studies.

We also identified phase-specific biomarker genes that allowed for phase-specific stratification of transcriptomic profiles regardless of severity. The acute phase (T1) - the most characteristic and readily distinguishable phase - provided the highest number of potential biomarker genes (770), as we anticipated after observing the distribution of groups in the consensus profile clustering analysis. This situation may be explained by the initial trauma that causes the rupture of cell membranes, axons, and blood vessels, generating a cascade of events in which many signals initiate stress response mechanisms and inflammatory response (*1*, *27*). Activating injury containment mechanisms in the acute phase results in the up-regulation of specific genes that return to their basal expression levels in subsequent phases (such as Il33 or Il6), evidencing the early invasion of non-neuronal cells responding to the injury stimuli (*52*). Similarly, genes related to the cell cycle, DNA replication, and DNA repair also exhibited this pattern (e.g., Rpa2, Hells, and Psrc1). This observation is further supported by functional enrichment analysis, where pathways associated with these categories become notably enriched in T1 but not in subsequent phases. The previously described activation of cell cycle genes after SCI can be explained by the proliferation/activation of mitotic cells (such as microglia or astrocytes) and the activation of apoptotic processes in damaged cells (*23*, *30*). In fact, it has been reported that inhibition of cell cycle gene expression may improve motor and cognitive recovery (*53–56*). Subacute phase-specific genes, including complement-related genes (e.g., Cfp, C1qa, C1qb, and C1qc), exhibited a distinct gene expression pattern peaking at T2. This pattern aligns with the dynamic behavior of cells such as macrophages and microglia that express and secrete complement proteins (*57*, *58*). Monocytes migrate to injury sites and differentiate into macrophages at approximately 3 dpi, culminating in a peak response at around 7 dpi while resident microglia reach an activation peak 1 week post-injury (*59*, *60*). Cfp exhibits a similar pattern in mice, and although this gene (along with Vav1 and Slc7a7) effectively distinguishes T2 in both rat and rat-mouse data, species-specific genes for T2 appear to exist in mice. Consistent up-regulation over time characterizes genes specific to T4 (peaking in the chronic phase), effectively distinguishing this phase from the others. Utilizing a combination of phase-specific genes supports the accurate classification of transcriptomic profiles for meta-analysis data and new transcriptomic profiles, such as those in the GSE218088 dataset. Additionally, while some of these genes may display species-specificity, common genes (such as Fosl1, Il6, Cfp, Sipa1, Cp, and Dbp) may find use in the temporal classification of rat and mouse samples. Experimental validation of Il6, Vcam1, and Vegfa confirmed the expression patterns observed in the meta-analysis and validated their similarity in the mouse dataset, further strengthening the robustness of our results. Thus, our study identified phase-specific biomarkers that may aid in selecting therapies based on transcriptomic profiles and improve patient stratification.

Our work provides a functional framework for assessing the degree of dysregulation in biological functions of interest. Our findings align with the findings of previous SCI research, characterized by a maintained up-regulation of functions related to the immune system and down-regulation of processes associated with the nervous system function and development. This phenomenon is influenced by other processes that sustain an unfavorable environment for functional recoveries, such as disturbed ionic homeostasis, vascular injury, ischemia, free radical stress, cell death (*19*, *24*, *27*), mitochondrial dysfunction (*31*), metabolic alterations (including the down-regulation of lipid and cholesterol metabolism) (*15*), or extracellular matrix remodeling (*61*). Considering this complex context, any strategy for developing effective therapies should include a combination of multiple targets, considering the interconnected nature of these systems. At the functional level, we observed differences between phases, with the decreasing significance of pathways related to cell cycle and RNA transcription as time progresses post-injury particularly noteworthy. However, in contrast to gene level results, we did not identify substantial differences between severities. Likewise, the constructed co-expression networks, comprising different genes with altered expression based on severity, demonstrated similarities in the enriched functions’ quantity and nature. This analysis corroborates the functional interconnection of genes with similar expression patterns (*62*, *63*) and agrees with the study of De Biase et al. (*64*), which failed to observe consistent alterations in functional patterns associated with injury severity, even given the apparent variations occurring at the gene expression level. This phenomenon may be explained by the design of functional enrichment techniques that capture the coordinated action of gene groups. Consequently, even though specific genes may display more pronounced dysregulation in severe injuries, their collective direction points toward the same outcome.

Our study also provides a way to infer the activity of master transcriptional regulators represented by pioneering evaluation of TF activity based on the expression levels of target genes, giving a new perspective for more efficient therapeutic interventions. Similar to the GSA outcomes, we identified temporal differences, with a higher number of TFs exhibiting differential activity in the initial phases that decreased over time. Additionally, a core group of TFs became permanently dysregulated. While we did not observe overall discrepancies in severity, the analysis unveiled three TFs with distinctive activity in severe injuries: Bhlhe22, Sp3, and Runx2. Bhlhe22 belongs to the bHLH family, known for its regulatory role in nervous system development. Specifically, Bhlhe22 functions by repressing the expression of target genes to facilitate accurate neuronal specification (*65–67*). Since its target genes showed to be up-regulated in severe injuries, it evidenced a deactivation of Bhlhe22 correlating with loss of neuronal identity features, which could explain the heightened impact on processes related to nervous system function and development in severe injuries. Previous studies indicate that both Sp3 and Sp1 display a significant increase in transcriptional activity during neuronal apoptosis, implying that their inhibition safeguards neurons from cell death (*68*). Our findings align with this trend, revealing a general increase in Sp1 activity and an elevation in Sp3 activity exclusively in moderate injuries. Runx2 functions in osteoblast differentiation and becomes up-regulated in Schwann cells following peripheral nerve injury (*69*); however, the role of Runx2 in SCI remains poorly understood, making this TF a potentially exciting target for further study.

In addition, we experimentally validated four TFs. Hes1 - a transcriptional repressor involved in neurogenesis (*70*) – displayed overexpression in our meta-analysis and the experimental validation and differential activation in all experimental groups. Previous studies support these findings, demonstrating Hes1 overexpression after SCI (*71*) and emphasizing the role of Hes1 in promoting neurogenesis through inhibition of its target genes, consequently enhancing functional recovery (*72*). Onecut1 – a transcriptional activator regulating the production, diversification, distribution, and maintenance of various neuronal populations (*73–75*) – became down-regulated following SCI; while we experimentally validated this result, the results were not wholly consistent with the expression pattern obtained from the meta-analysis, possible by an inter-experimental deviations. Activity inference analysis indicated deactivation, especially in the early phases of injury. These findings suggest overexpression or increased activity levels could benefit neuronal development and function. Our bioinformatic and experimental results also indicated an up-regulation of Cebpd - a transcriptional activator that regulates genes involved in immune responses (*76*) - after injury, accompanied by differential activation in acute phases. Previous studies reported Cebpd overexpression following injury (*76*, *77*), along with enhanced expression during functional recovery in mice deficient in this gene (*77*). Jun - a transcriptional activator highly induced following neuronal damage (*78*, *79*) – became consistently overexpressed and activated in all experimental groups; in a related study, Zhang et al. proposed that miR-152 overexpression inhibited inflammatory responses and promoted functional recovery by inhibiting Jun (*79*).

Of a great interest is the correlation we found within the expression profiles in human samples and the rat spinal cord transcriptional data. We identified a 12-gene signature predicting injury severity in human blood and rat spinal cord data. Among these 12 genes, MMP9 has been proposed as a potential severity biomarker in species including humans, rats, and dogs (*46*, *48*, *50*)). The clinical application of this signature could be valuable, enabling patient stratification through simple techniques such as blood extraction and transcriptional analysis. Although the ASIA system, accompanied by the anatomical evidence of the extent of the affected and preserved tissue by MRI, represents the standard for predicting SCI severity, there exists a need to find new operative and non-invasive methods for its prognosis. The ASIA system relies on neurological status evaluations through sensory and motor exams; however, this method suffers from limitations (*40*). Additionally, the ASIA classification does not consider patient heterogeneity; for example, younger individuals may suffer from enhanced recovery potential (*80*). The 12-gene signature could also aid laboratory research using rat models. Currently, assessing functional improvements in response to treatment requires locomotion tests or post-mortem analysis. Evaluating treatment efficacy through in vivo blood extraction would simplify progress monitoring; however, further investigation will be required to explore the prognostic capacity of these genes in detail with larger sample sizes in humans and additional relevant models.

The development of the Meta-SCI app during this study offers an additional dimension. This platform extends beyond a mere results repository; Meta-SCI serves as an analysis tool for the research community. The app allows researchers to access the different analyses and explore gene-function relationships that match their research interests. Also, Meta-SCI includes two injury models, allowing researchers to choose that which best suits their needs. Consequently, the Meta-SCI app emerges as a valuable resource for hypothesis generation and validation and constitutes a transcriptional reference database for studying SCI.

Our study offers a comprehensive systems biology frame of SCI, providing a holistic view of the gene and functional levels and the dimensions of severity and phase. We identified potential biomarker genes that effectively stratified transcriptomic profiles based on time and severity, offering critical insights into SCI progression. Thanks to our user-friendly Meta-SCI web application, researchers can explore this extensive results repository, tailoring their investigations to their fields of interest. Our study sheds light on the intricate mechanisms underlying SCI and equips researchers with the means to translate these insights into diagnostic and therapeutic interventions.

## Materials and Methods

### Study search and selection

A systematic review and selection of studies were conducted in September of 2022. The search was performed in two public databases for transcriptomic data - GEO and ArrayExpress - according to PRISMA statement guidelines (*81*) and carried out with the keywords “spinal cord injury.” The inclusion criteria were: (1) organism, Rattus norvegicus; (2) tissue, thoracic spinal cord; and (3) RNA sequencing or microarray data from Affymetrix or Agilent used as gene expression platforms. The exclusion criteria were: (1) studies without control samples (sham or naive rats); (2) individual samples from rats that had received any type of treatment; and (3) samples rostral or caudal to the epicenter of the injury.

### Data preprocessing

For microarray studies, normalized data were downloaded with the GEOquery package and probes were annotated to their corresponding gene symbol using platform-specific annotation packages. For RNA-seq studies, raw count matrices were manually downloaded from GEO and identifiers were converted to gene symbols using the org.Rn.eg.db package. In all cases, the median of the expression values of duplicated identifiers was calculated.

### Exploratory analysis

An exploratory analysis was performed (including boxplot, PCA, and clustering analysis) to observe sample distribution in the groups of interest and evaluate anomalous behavior and possible batch effects. Samples from the GSE183591 study were taken in two different series, separated into two blocks in the PCA. This batch effect was taken into consideration for DGE analysis. The negative values detected in boxplots for the GSE2599 and GSE464 datasets were normalized by adding the minimum of all values followed by a logarithmic transformation of the data.

### Differential gene expression analysis

Each injury group was compared with its corresponding control within the study to identify DEGs. DGE analysis of microarray platforms was carried out using the limma package (*82*). To ensure comparable results between technologies, RNA-seq datasets were analyzed using the limma-voom pipeline, which involves the removal of genes with low counts and a logarithmic transformation of the expression matrix. p-values were corrected with the Benjamini–Hochberg (BH) method (*83*).

### Gene expression meta-analysis

DGE results were integrated to obtain a transcriptional consensus signature for each comparison. A meta-analysis was performed for each gene in at least two studies in each comparison. The random-effects model proposed by Der Simonian & Laird (*84*), implemented in the metafor package (*85*), was used to evaluate the combined effect. This method weighs the heterogeneity of each study and incorporates their inherent variability into the overall estimate of the measure of effect. Consequently, studies with greater variability will have lower weight in this measure. In this case, logFC was used to measure the effect, while the variance was used to measure variability. Thus, values for the p-value, logFC, and 95% confidence interval (CI) were calculated for each gene evaluated in the meta-analysis. Since multiple meta-analyses were performed, p-values were adjusted using the BH method. Adjusted p-values lower than 0.1 were considered significant. Funnel and forest plots were used to assess the variability and measure each study’s contribution to the meta-analysis.

### Clustering analysis

All clustering analyses were performed on the gene consensus signature matrix. We constructed this matrix with consensus logFC values from the meta-analysis, selecting genes evaluated in all groups and significant in at least one group. The mixOmics package for PCA was used (*86*). For hierarchical clustering analysis, the Euclidean distance was first calculated as input for the hclust function.

### Biomarker gene identification

In this context, biomarker genes were defined as those displaying more significant deregulation in certain groups of interest than others, allowing the stratification of different groups based on their expression pattern. A limma test was applied to identify possible biomarker genes that could classify samples according to severity/phase. he four severe were compared against the four moderate groups of samples to elucidate severity-specific biomarker genes. Phase-specific biomarker genes common to both severities and specific to moderate or severe injury were also explored. Detailed comparisons are listed in table S4. Subsequently, the top 10 genes with the lowest p-values from each comparison were selected to assess their classification ability via clustering analysis comprising PCA and hierarchical clustering.

### Gene set analysis

A functional enrichment analysis was performed for each gene consensus signature with the GSA method implemented in the mdgsa R package (*87*). A gene ranking and functional annotation are required as input for GSA. The ranking is made by ordering all genes according to the p-value obtained in meta-analysis and the logFC value sign. For functional enrichment annotation, three annotation databases have been included: the biological processes of Gene Ontology (*88*), KEGG pathways (*89*), and Reactome pathways (*26*). Gene sets with less than 10 or more than 500 genes were excluded. P-values were corrected with the BH method.

### Transcription factors activities

The identification of differentially active TFs used the msviper function of the viper package (*90*) with each of the gene consensus signatures as input. Mouse regulons from the DoRothEA package (*91*) with a confidence level of A, B, C, or D were selected, excluding those with less than 10 genes (table S11). The p-values were corrected with the BH method.

### Construction of gene co-expression networks

First, in order to find genes with similar expression patterns we used the clust software (*92*) with the following arguments: -n 0 -t 100 -cs 4. Gene consensus signatures matrix from moderate and severe injuries were used separately as input. A PPI was then constructed for each cluster of co-expressed genes obtained from clust software. The STRINGdb package with the interactions of STRING version 11.5 (*93*) and medium confidence of 0.400 were used to build PPI networks. The PPI network of each cluster was divided into different subnetworks of more closely related genes using the fastgreedy algorithm implemented in the get_clusters function.

Each subnetwork was then classified into manually pre-defined functional categories of interest. The get_enrichment function was first used to obtain overrepresented functional terms in each subnetwork. Subsequently, the biological processes were selected from the Gene Ontology, KEGG, and Reactome pathways, ad hoc classifying them into pre-established general categories. Finally, the frequency distribution of each category was calculated for each subnetwork, assigning the category with the highest frequency as that which best defines the network.

### Bioinformatic validation datasets

For the mouse transcriptomic dataset, a normalized count matrix provided from the supplementary material provided by Li et al. (*36*) (https://doi.org/10.6084/m9.figshare.17702045) was downloaded. Samples representative of the four temporal phases established in our study (1, 7, 28, and 42 dpi) and controls were selected (table S19). For the GSE218088 dataset, the normalized expression matrix was downloaded using GEOquery, and the probes were annotated to the gene symbol. Those genes that displayed significance in at least one comparison and were present in all groups of our meta-analysis were selected for both datasets. A DGE analysis using the limma package was performed, comparing each injured group against uninjured controls.

### Experimental validation (qPCR)

A severe traumatic SCI model in adult female Sprague Dawley rats was employed for RNA isolation and qPCR in silico data validation. The animals were housed at the Animal Experimentation Unit of the Research Institute Príncipe Felipe (Valencia, Spain) under standard conditions. All experimental procedures adhered to the guidelines established by the European Communities Council Directive (86/609/ECC), the Spanish Royal Decree 53/2013, and the Animal Care and Use Committee of the Research Institute Príncipe Felipe (2021/ VSC/PEA/0032).

Rats were subcutaneously pre-medicated with morphine (2.5 mg/kg) and anesthetized with 2% isoflurane in a continuous oxygen flow of 1 L/min. Laminectomy was performed on thoracic vertebrae 8–9 to expose the spinal cord, and severe SCI was induced at the thoracic vertebrae 8 level by contusion, applying a force of 250 kdyn using the Infinite Horizon Impactor, as previously described (*13*, *94*). Post-surgery care included manual bladder drainage twice a day until vesical reflex recovery and subcutaneous administration of 5 mg/kg of Enrofloxacin (Alsir) for 7 days, as well as 0.1 mg/kg of Buprenorphine twice a day for 4 days after each intervention.

At each endpoint: 1 dpi (n = 3), 7 dpi (n = 3), 28 dpi (n = 4), and 56 dpi (n = 4) after injury to represent (T1, T2, T4, and T8, respectively), animals were overdosed of sodium pentobarbital (100 mg/kg) and transcardially perfused with a 0.9% saline solution. The spinal cord tissue was extracted, and the injury epicenter was immediately frozen in liquid nitrogen and stored at – 80 °C until use. RNA extraction was performed using the TriZol standard method, followed by an additional cleanup step using RNeasy MinElute Cleanup (Qiagen, Germany) to ensure sample quality (A260/280 ≈ 2 and A260/230 ≥ 1.8). Reverse transcription was performed using the high-capacity RNA-to-cDNA™ kit (Applied Biosystems, Massachusetts, USA).

Specific primers for each gene of interest were designed using primer-BLAST (NCBI, Maryland, USA) and validated by efficiency curve performance. qPCR was conducted in triplicate using AceQ SYBR qPCR Master Mix (ThermoFisher) in the Light-Cycler 480 detection System (Roche, Basel, Switzerland). Ct data were calculated with the LightCycler 480 relative quantification software (Roche, Basel, Switzerland). GAPDH mRNA levels served as an internal control for normalization.

### Analysis of human blood samples

The GSE151371 normalized gene expression matrix was downloaded from GEO, and the expression values were log-transformed. The samples corresponding to the AIS A, AIS D, and the healthy control groups were then selected for further analysis (table S20). DGE analysis was then performed to compare AIS A vs. AIS D with the limma package following the same pipeline as the previous analyses. Finally, significantly altered genes (FDR < 0.1) were selected to compare with the list of rat biomarker genes. The list of intersecting genes was used for the clustering analysis of both biological systems. The healthy controls from the GSE151371 dataset were also included for visualization and clustering analysis.

### Meta-SCI app

Meta-SCI app is powered by R and RStudio Shiny. This straightforward design ensures easy access and efficient analysis for users. Meta-SCI is deployed on a shinyapps.io server, available at https://metasci-cbl.shinyapps.io/metaSCI/. Plots are generated using ggplot, plotly, and heatmaply. All data processing and analysis are carried out using R.

## Acknowledgments

The authors thank the Principe Felipe Research Center (CIPF) for providing access to the cluster, co-funded by European Regional Development Funds (FEDER) in Valencian Community 2014-2020. The authors also thank Stuart P. Atkinson for reviewing the manuscript.

## Author contributions

Conceptualization: FGG

Methodology: RGR, MRH, BMR, VMM, FGG

Investigation: FGG, MRH, BMR, VMM, RGR

Visualization: RGR, MRH

Supervision: MRH, VMM, FGG

Writing—original draft: RGR, MRH, BMR, VMM, FGG

Writing—review & editing: RGR, MRH, BMR, VMM, FGG

## Data and materials availability

The large volumes of data and results generated in this study are freely available through the Meta-SCI app: https://metasci-cbl.shinyapps.io/metaSCI/. This study analyzed transcriptomic data available in the Gene Expression Omnibus database with accession numbers: GSE183591, GSE182796, GSE138637, GSE133093, GSE115067, GSE102964, GSE104317, GSE93249, GSE52763, GSE45006, GSE46988, GSE29488, GSE2599, GSE464, GSE151371 and GSE218088.

